# Regulation of *Chlamydia* RsbU phosphatase activity by the enolase product, phosphoenolpyruvate

**DOI:** 10.1101/2022.05.13.491914

**Authors:** Christopher Rosario, Ming Tan

## Abstract

The intracellular pathogen *Chlamydia* temporally regulates the expression of its genes but the upstream signals that control transcription are not known. The best studied regulatory pathway is a partner switching mechanism that involves an anti-sigma factor RsbW, which inhibits transcription by binding and sequestering the sigma subunit of RNA polymerase. RsbW is itself regulated by an anti-anti-sigma factor RsbV whose phosyphorylation state is controlled by the phosphatase RsbU. In this study, we showed that *Chlamydia trachomatis* RsbU requires manganese or magnesium as a cofactor and dephosphorylates RsbV1 and RsbV2, which are the two chlamydial paralogs of RsbV. The gene for RsbU is adjacent to the enolase gene in a number of *Chlamydia* genomes, and we showed that *eno* and *rsbU* are co-transcribed from the same operon. In other bacteria, there is no known functional connection between the Rsb pathway and enolase, which is an enzyme in the glycolytic pathway. We found, however, that *Chlamydia* RsbU phosphatase activity was inhibited by phosphoenolpyruvate (PEP), the product of the enolase reaction, but not by 2-phosphoglycerate (2PGA), which is the substrate. These findings suggest that the enolase reaction, and more generally glucose metabolism, may provide an upstream signal that regulates transcription in *Chlamydia* through the RsbW pathway.

**IMPORTANCE:** The RsbW pathway is a phosphorelay that regulates gene expression in *Chlamydia* but its upsteam signal has not been identified. We showed that RsbU, a phosphatase in this pathway is inhibited by phosphoenolpyruvate, which is the product of the enolase reaction. As enolase is an enzyme in the glycolytic pathway, these results reveal an unrecognized link between glucose metabolism and gene regulation in chlamydiae. Moreover, as these intracellular bacteria acquire gluose from the infected host cell, our findings suggest that glucose availability may be an external signal that controls chlamydial gene expression.

## INTRODUCTION

*Chlamydia* is a pathogenic bacterium that requires a host cell for growth and replication. The intracellular infection is characterized by an unusual developmental cycle in which the bacterium converts between two forms, an infectious form called an elementary body (EB) and a noninfectious but metabolically active form called a reticulate body (RB) (Abdelrahman and Belland, 2005; Moulder, 1991). During the early stage of the infection, the EB enters the host cell, forms a membranous vacuole called an inclusion, and converts into an RB. RBs then divide repeatedly during the midcycle stage of the infection. Finally at the late stage of the developmental cycle, RBs convert asynchronously into EBs, and then exit the host cell via lysis or extrusion by 48-72 hours post infection (hpi) (Hybiske and Stephens, 2007).

Chlamydial genes are transcribed in three temporal waves that correspond to these three developmental stages (Belland et al., 2003; Nicholson et al., 2003; Rosario et al., 2020; Shaw et al., 2000). For example, early genes are transcribed within the first few hours of EB entry into the host cell (Belland et al., 2003; Hayward et al., 2021; Humphrys et al., 2013). Midcycle genes are transcribed during RB replication and are proposed to be regulated by changes in chlamydial DNA supercoiling (Niehus et al., 2008; Orillard and Tan, 2015). Late genes are transcribed at the time of RB-to-EB conversion and are regulated by a transcription factor EUO (Rosario et al., 2014; Rosario and Tan, 2012). In addition, a subset of late genes are transcribed by an alternative form of RNA polymerase containing sigma28 (σ^28^) instead of the major sigma factor, sigma66 (σ^66^) (Yu et al., 2006a; Yu et al., 2006b; Yu and Tan, 2003).

In addition to these mechanisms of gene regulation, *Chlamydia* may control RNA polymerase activity through an anti-sigma factor that binds and sequesters the sigma factor (Rosario et al., 2020). The sigma factor is the subunit of RNA polymerase that allows it to recognize and bind specific promoter DNA sequences and thereby transcribe its target genes. *Chlamydia* encodes an anti-sigma factor RsbW, which binds the sigma factor sigB (σ^B^) in *Bacillus* (Helmann, 1999). There is no σ^B^ ortholog in *Chlamydia*, but chlamydial RsbW has been proposed to bind and inhibit either σ^66^ or σ^28^, which are two of the three chlamydial sigma factors (Douglas and Hatch, 2006; Hua et al., 2006; Thompson et al., 2015).

*Chlamydia* also contains components of a signaling pathway that regulates RsbW and σ^B^ in *Bacillus* (Figure 1) (Hua et al., 2006). The core of this pathway is a partner switching mechanism in which RsbW binds either its sigma factor or an anti-anti-sigma factor RsbV, depending on the phosphorylation state of RsbV. Phosphorylated RsbV cannot bind RsbW, allowing this anti-sigma factor to bind and inhibit its cognate sigma factor. However, when RsbV is unphosphorylated, it binds RsbW, which frees up the sigma factor and activates transcription.

**Figure 1.**
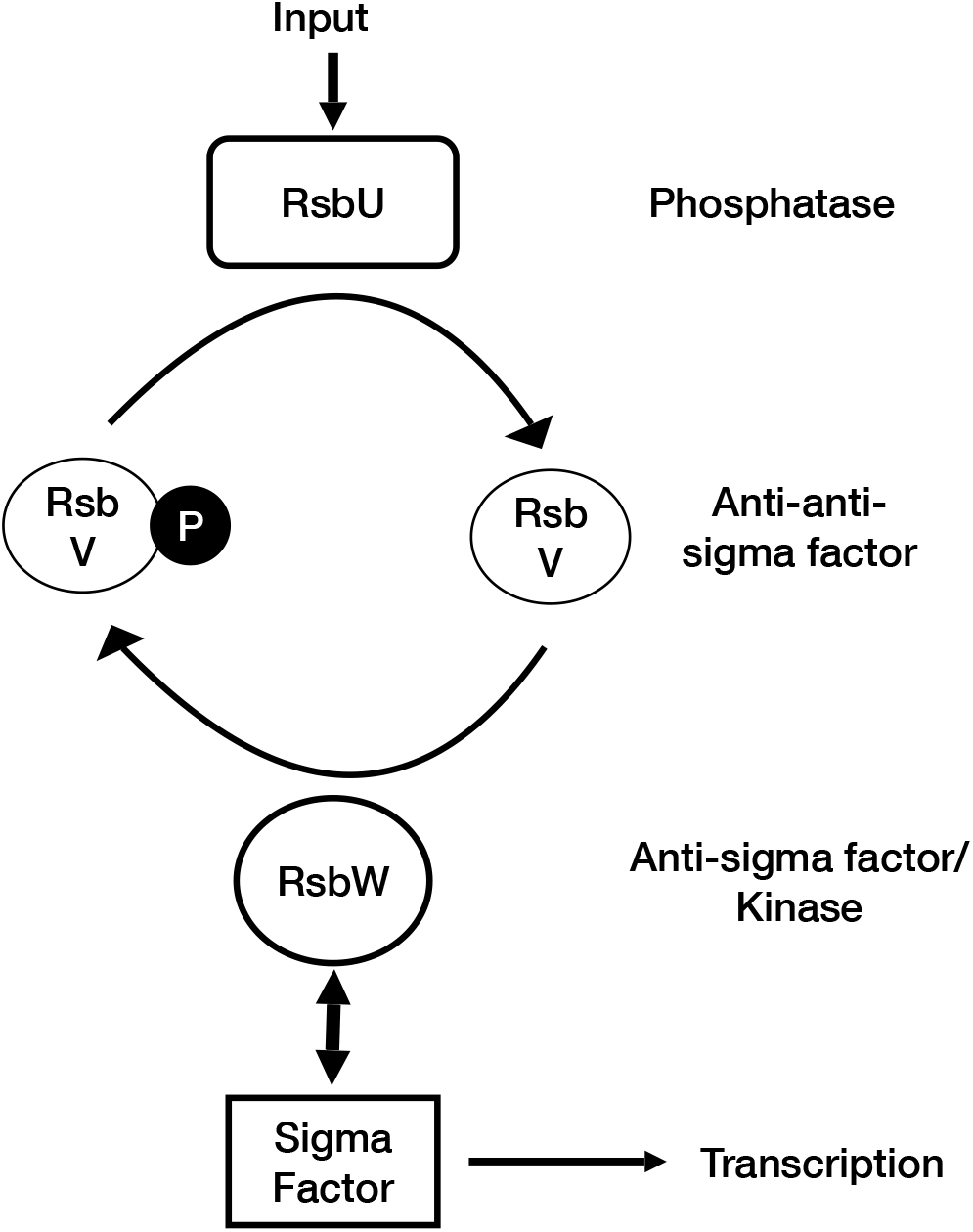
Diagram of the RsbW partner switching mechanism. RsbW binds to either the sigma factor or the anti-anti-sigma factor RsbV, depending on the phosphorylation state of RsbV. RsbW binds to unphosphorylated RsbV but not phospho-RsbV. The balance between RsbV and phospho-RsbV is determined by the kinase activity of RsbW and the phosphatase activity of RsbU. RsbU is thus a positive regulator of transcription that allows RsbV to bind and sequester RsbW, which frees up the sigma factor for transcription of target genes.

RsbU is the phosphatase that dephosphorylates RsbV, and it thus serves as a positive regulator of transcription. *Chlamydia* has two RsbV paralogs, and *C. trachomatis* RsbU has been shown to dephosphorylate RsbV1 but has not been tested on RsbV2 (Thompson et al., 2015). Soules and Hefty recently showed that intermediates of the TCA cycle bound to the periplasmic domain of *C. trachomatis* RsbU, which is separate from its phosphatase domain (Soules et al., 2020). However, it is not known if chlamydial RsbU phosphatase activity can be regulated by these TCA intermediates or other factors, and thus the upstream signal that controls the Rsb pathway in *Chlamydia* has not been identified.

In this study, we investigated how the phosphatase activity of *C. trachomatis* RsbU is regulated. We were intrigued by the location of the enolase gene next to the RsbU gene in several *Chlamydia* spp. and explored if this glycolytic enzyme could somehow regulate RsbU. We found that phosphoenolpyruvate (PEP), which is the product of the enolase reaction, inhibited RsbU enzymatic activity. This finding suggests that the glycolytic pathway may have an unexpected role in regulating gene expression in *Chlamydia*.

## MATERIALS AND METHODS

### Plasmids and strains

Plasmids used in this assay are listed in Table 1. *C. trachomatis* LGV serovar L2 434/Bu was used as a source of genomic DNA for cloning and for RNA extraction. L929 mouse fibroblast cells were used for *C. trachomatis* infections. Cells and infections were grown in RPMI 1640 supplemented with 25 mM HEPES and 5% FBS. *E. coli* BL21 was used as a source of genomic DNA for cloning and for protein purification.

### Protein purification

For protein expression, *C. trachomatis* or *E. coli* genes were cloned into pRSET C (Thermo Fisher), which adds a 6x His moiety to the N-terminus of the recombinant protein. Plasmids were used to transform *E. coli* strain BL21, and transformed cells were grown in LB broth. Protein expression was induced with 1 mM IPTG for 2 hr. Cells were pelleted and resuspended in buffer N (10 mM Tris HCl pH 8.0, 300 mM NaCl, 1 mM MgCl_2_, and 10 mM 2-mercapoethanol) containing 25 mM imidazole. Cells were lysed by sonication in a Bradford digital sonifier 250D for 2 minutes at 25% output. Cell debris was pelleted and the supernatant was incubated with a 2 ml slurry of Ni-NTA (Sigma) for 1 hr at 4°C with agitation. Bound protein was washed with 50 ml buffer N containing 25 mM imidazole. Protein was eluted with buffer N containing 250 mM imidazole. Eluted protein was dialyzed against storage buffer (10 mM Tris-HCl [pH 8.0], 100 mM NaCl, 10 mM 2-mercaptoethanol, 30% [v/v] glycerol). Dialyzed protein was aliquoted and stored at −70°C.

### RsbU phosphatase assay

Purified recombinant RsbV1 or RsbV2 (20 ug) was radiolabeled by incubating with purified recombinant RsbW (5 ug) and approximately 30 uCi [gamma-^32^P]-ATP (10 mCi mmol-1; Perkin Elmer) in Cutsmart buffer (NEB) at 37°C for 30 min. Radiolabeled protein was separated from free ^32^P-ATP by centrifugation in mini Quick Spin DNA column (Roche). Radiolabeled protein was diluted in glycerol (20% [v/v] final concentration) and stored at −20°C.

For the phosphatase assay, radiolabeled RsbV protein (approximately 0.02 μg; 5 μM) was incubated with purified recombinant RsbU (0.5 μg; 6 μM) in buffer P (10 mM Tris, pH 7.5) with magnesium chloride or manganese chloride (as indicated in text) for 30 min at 37°C. For assays with other ions, Mg was not present in buffer P. Reactions were stopped by addition of SDS-PAGE loading buffer. Samples were analyzed by SDS-PAGE electrophoresis on a 18% SDS-PAGE gel. Gels were exposed to a phosphoimager screen, which was then scanned on an Amersham Typhoon Biomolecular imager.

### RT-PCR

L929 mouse fibroblast cells were infected with *C. trachomatis* L2 at an MOI of 3. At indicated times, infected cells were collected with glass beads in PBS buffer. Cells were pelleted and used for RNA extraction using a Macherary-Nagel RNA extraction kit. RNA was treated with RQ1 DNAse (Promega) to remove any DNA. Approximately 5 μg total RNA was used for reverse transcription with MMLV reverse transcriptase, with a primer located 1697 bp downstream of the ATG translational start site of the *rsbU* gene. cDNA was used for PCR using primers amplifying a 1309 bp fragment located 1162 bp downstream (5’-GGGTCTCTTTCCCGTTCTGA) of the ATG translational start site of the *eno* gene to 1057 bp downstream (5’-GGATGGTGTTGATTGCGACG) of the ATG translational start site of the *rsbU* gene. PCR was performed with Bioneer AccuPower *Pfu* PCR premix with these amplification conditions: 30 sec at 95°C, 45 sec at 57°C, 90 sec at 71°C for 40 cycles. PCR products were analyzed by electrophoresis on a 1% agarose gel in TBE. Gels were stained with ethidium bromide, and bands were visualized by UV light with an Amersham Imager 680.

### 5’ RACE

First Choice RLM-RACE reactions (Thermo Fisher) were performed with RNA extracted from infected L929 cells at 18 and 32 hpi. 5’ RACE assays were performed per manufacturer’s instructions. A primer (5’-GCAGGGTGAGGAGGAACTTC) located 1697 bp downstream of the ATG translation start site of *rsbU* was used for cDNA synthesis. The same primer was used to amplify a PCR product to the modified 5’ end of the mRNA. Based on published transcription start sites (Albrecht et al., 2010), these primers generate expected PCR products of ~3.1 kB and a ~1.6 kB for the *eno* and *rsbU* promoters, respectively. PCR products were analyzed by electrophoresis on a 1% agarose gel in TBE. Gels were stained with ethidium bromide and bands were visualized by UV light with an Amersham Imager 680.

### Transcription assays

Transcription assays were performed as previously described (Yu et al., 2006a). Essentially, 2.5 μM recombinant EUO was incubated with 13 nM plasmid DNA at room temperature for 15 minutes. Transcription was initiated with 0.4 U *E. coli* holoenzyme (Epicentre) in the presence of ^32^P-CTP at 37°C for 15 min. Transcripts were resolved on an 8 M urea-6% polyacrylamide gel. Gels were placed on Whatman paper and exposed to a phosphorimager screen. The phosphoimager screen was scanned on an Amersham Typhoon Biomolecular imager. Band intensities were quantified using Quantity One software (Bio-Rad). For each promoter, the relative transcript levels were calculated by measuring the transcript levels in the presence of EUO and normalizing to levels in the absence of EUO. Values are reported as the mean of the relative transcript levels with standard deviations from at least three individual experiments.

### Enolase assay

Enolase assays were performed using an enolase activity kit (Sigma MAK178). Essentially, purified recombinant His-tagged enolase was incubated in 2-phosphoglycerate in a Costar 3603 clear, flat bottom 96-well assay plate. The production of phosphoenolpyruvate is proportional to the amount of hydrogen peroxide produced. The production of hydrogen peroxidase was measured by fluorometrically (λ_ex_= 535 nm/λ_em_ = 587 nm) on a SpectraMax i3x, with readings taken every 5 minutes for 30 minutes at 25°C. Activity was calculated as the rate of nmol of hydrogen peroxide produced per time (minutes) per mL of sample.

## RESULTS

### Co-factor requirement of *C. trachomatis* RsbU

We first determined the metal co-factor that is necessary for the phosphatase activity of chlamydial RsbU. RsbU belongs to the PP2C phosphatase family that requires magnesium or manganese for activity, but is inhibited by zinc (Das et al., 1996). We were unable to purify full-length recombinant RsbU, which is a large transmembrane protein, but successfully purified a truncated form of RsbU containing the C-terminal 260 amino acids, which encompasses the phosphatase domain but lacks the periplasmic domain. We then tested our RsbU polypeptide in an *in vitro* phosphatase assay that utilized recombinant *C. trachomatis* RsbV1 and RsbV2 that had been phosphorylated by RsbW with ^32^P-radiolabel.

RsbU dephosphorylated RsbV1 and RsbV2, but there were differences in its activity against these two substrates (Figure 2). RsbV1 was dephosphorylated by RsbU in the presence of either Mn^2+^ or Mg^2+^, but RsbV2 was only dephosphorylated at high Mn^2+^ concentration and not with Mg^2+^ (Figure 2A). The concentration of Mn^2+^ required for comparable RsbU phosphatase activity against RsbV1 was 100-fold lower than for Mg^2+^ (Figure 2A, lanes 4 and 5), which suggests that Mn^2+^ is a better co-factor than Mg^2+^. There was no phosphatase activity with Ca^2+^ or Zn^2+^, and in fact Zn^2+^ inhibited RsbU activity against both RsbV1 and RsbV2 (Figure 2A, 2B). These results demonstrate that RsbU, like other PP2C phosphatases, can use manganese or magnesium as a co-factor while its activity is inhibited by zinc (Sajid et al., 2015).

**Figure 2.**
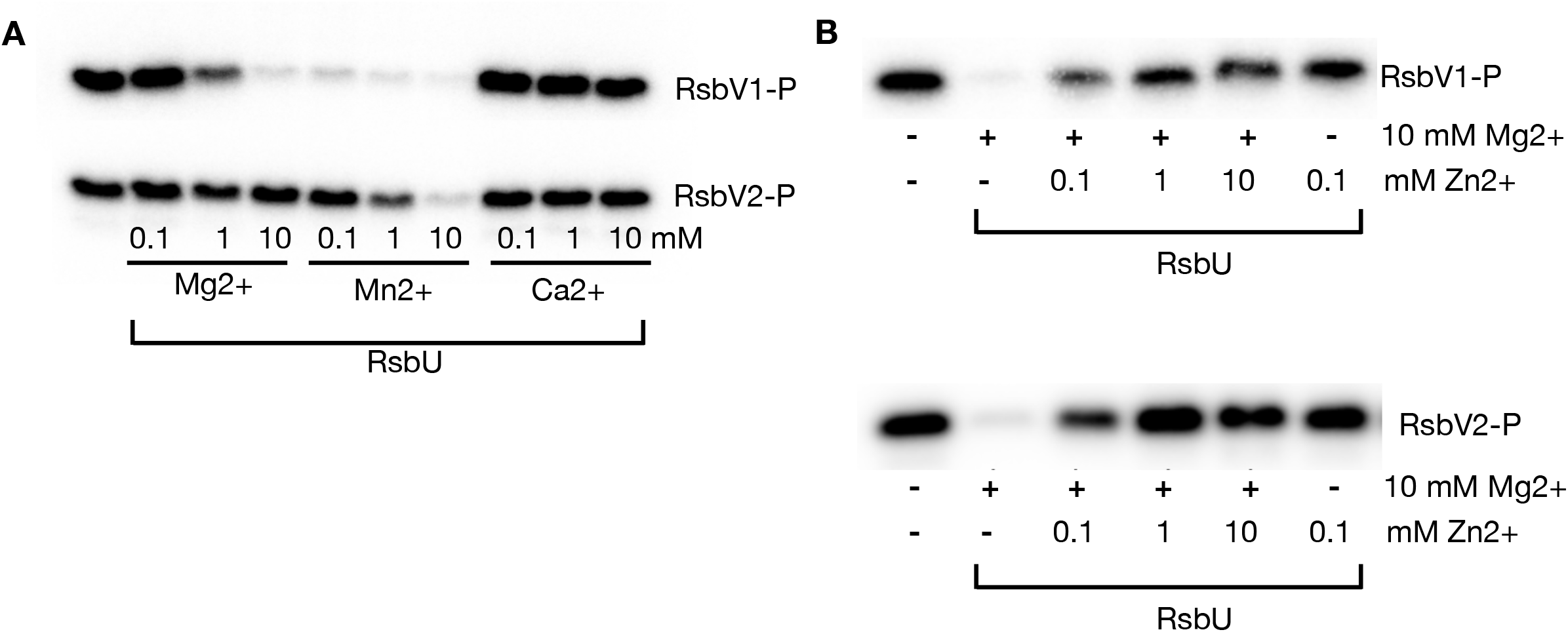
RsbU dephosporylates RsbV1 and RsbV2. A. *In vitro* RsbU phosphatase assay with purified recombinant RsbU (6 μM) and ^32^P-labelled RsbV1 or RsbV2 (5 μM each). Reactions were performed in the presence of increasing concentrations (0.1, 1, and 10 mM) of MgCl_2_, MnCl_2_, or CaCl_2_. Labeled RsbV was visualized by autoradiograph. B. Inhibition of RsbU phosphatase activity with increasing concentration (0.1, 1, and 10 mM) of ZnCl_2_ in the presence of co-factor (MgCl_2_ or MnCl_2_)

### Transcriptional organization of the RsbU and enolase genes

While searching for potential regulators of RsbU, we were intrigued by enolase, because the genes for these two proteins are adjacent in many *Chlamydia* genomes (Figure 3A) (Kanehisa, 2019; Kanehisa et al., 2021; Kanehisa and Goto, 2000). We first used RT-PCR to investigate if *eno* and *rsbU* are in the same operon. Using a 5’-primer in *eno* and a 3’-primer in *rsbU*, we successfully amplified a 1.3 kb PCR product from chlamydial RNA harvested from L929 mouse fibroblast cells infected with *C. trachomatis* serovar L2 (Figure 3B). These data provide evidence that *eno* and *rsbU* are co-transcribed on the same polycistronic message in this bacterium.

**Figure 3.**
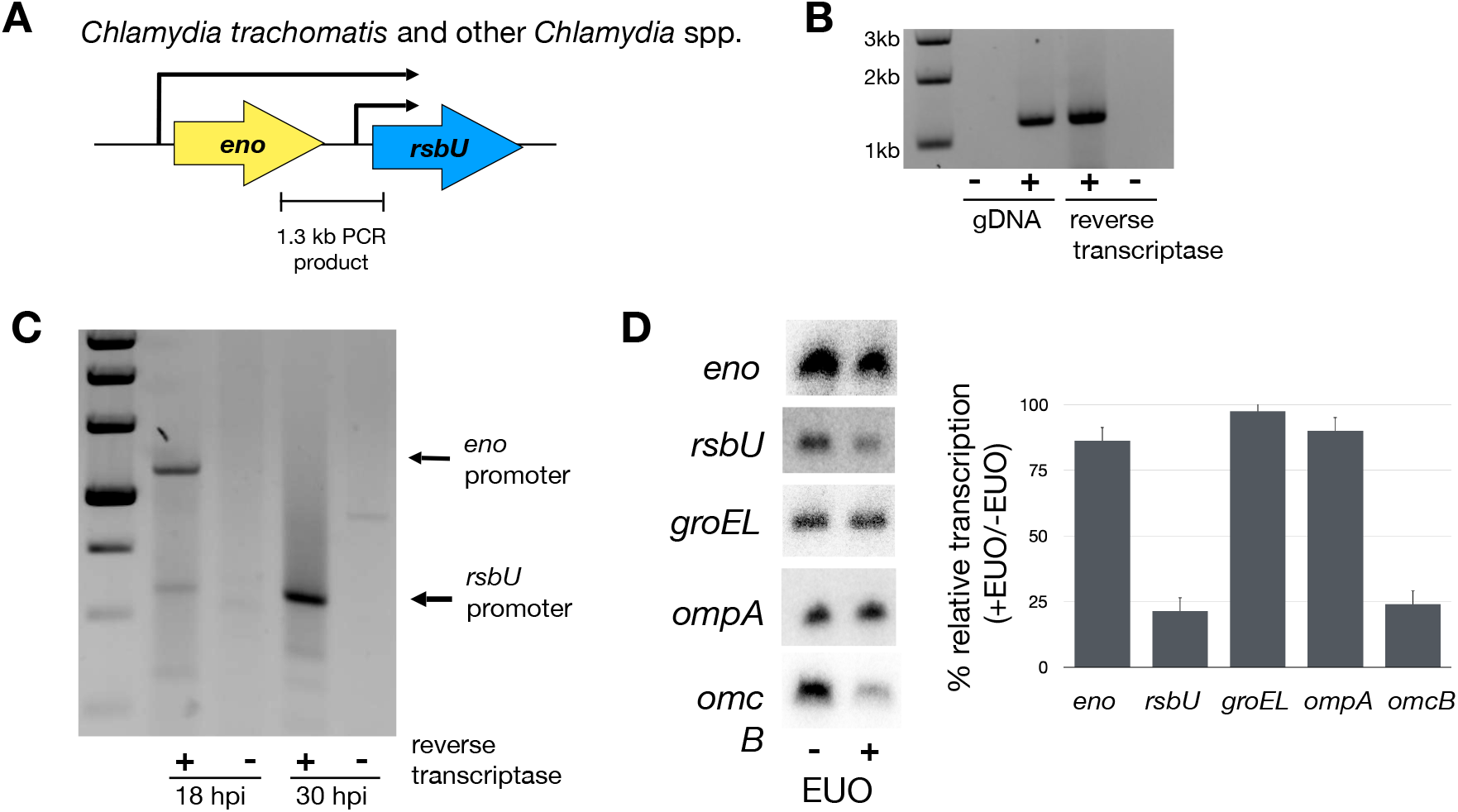
The *eno* and *rsbU* genes are in an operon. A. Gene organization of *eno* and rsbU *genes* in *C. trachomatis* and other *Chlamydia* spp. Arrows above the genes represent transcriptional starts sites identified by RNA-seq (Albrecht et al., 2010). The 1.3 kB PCR product used for RT-PCR is indicated below the *C. trachomatis* gene diagram. B. 1.3 kb PCR product was detected by reverse transcription of *C. trachomatis* RNA collected at 24 hpi. Primers used in the PCR reaction amplified a segment that annealed to the 3’ end of *eno* and 5’ end of *rsbU. C. trachomatis* genomic DNA (gDNA) was used as a control. C. 5’ RACE showing differential expression of the *eno* and *rsbU* promoters. 5’ RACE was performed wth a primer that annealed to the 3’ end of *rsbU* and specific 5’ primers for the *eno* and *rsbU* promoters. *C. trachomatis* RNA was collected at 18 and 30 hpi. PCR products were visualized on a 1% agarose gel. Predicted PCR products corresponding to transcripts from the *eno* and *rsbU* promoters are indicated by the arrows. D. EUO-mediated repression of the *rsbU* but not *eno* promoter. *In vitro* transcription assays of the promoters of *eno*, *rsbU, omcB* (positive control), *groEL*, and *ompA* (negative controls) were transcribed with *E. coli* RNA polymerase in the presence or absence of 5 μM EUO. Transcripts were quantified with Quantity One software. For each promoter, relative transcription was calculated as the percent of transcripts in the presence of EUO to transcripts in the absence of EUO. Values are from the average of at least three independent experiments with standard deviation indicated by the error bar.

We then used 5’ RACE to examine the temporal transcription of *eno* and *rsbU* in *C. trachomatis*-infected cells from their published transcription start sites (Albrecht et al., 2010). At 18 hpi, we detected a 5’ RACE PCR product from the *eno* promoter and a weak product from the *rsbU* promoter (Figure 3C). In contrast, at 30 hpi, the 5’ RACE PCR product was primarily from the *rsbU* promoter with no product from the *eno* promoter. Together these findings suggest that *eno* and *rsbU* are co-transcribed from a promoter that is active at 18 hpi, which is midcycle in the chlamydial developmental cycle, but downregulated at 30 hpi, which is a late time. In addition, detection of *rsbU* transcripts at 30 hpi suggests that *rsbU* may have its own promoter that is preferentially transcribed at late times.

We performed *in vitro* transcription assays to study the promoters for *eno* and *rsbU*. Both promoters were transcribed by *E. coli* σ^70^ RNA polymerase (which is equivalent to *C. trachomatis* σ^66^ RNA polymerase in transcribing *Chlamydia* σ^66^ promoters) (Figure 3D) (Rosario and Tan, 2012). As *rsbU* appears to be a late gene, we investigated if it is regulated by the late regulator EUO (Rosario et al., 2014; Rosario and Tan, 2012). EUO repressed the *rsbU* promoter, and a positive control late promoter *omcB*, but not the *eno* promoter or two negative control midcycle promoters, *groEL* and *ompA* (Figure 3D). These findings provide evidence that *rsbU* can be expressed from two promoters, a midcycle promoter for the *eno-rsbU* operon and an internal promoter that transcribes *rsbU* at late times.

### RsbU phosphatase activity is regulated by the product of the enolase reaction

To study chlamydial enolase, we first checked if *C. trachomatis* enolase is enzmatically active. Enolase is an enzyme in the glycolytic pathway that catalyzes the conversion of 2PGA to PEP in bacteria and many other organisms (Figure 4A) (Kanehisa, 2019; Kanehisa et al., 2021; Kanehisa and Goto, 2000). Using an *in vitro* assay, we showed that purified recombinant *C. trachomatis* enolase converted 2PGA to PEP, in a similar manner as *E. coli* enolase (Figure 4B). In contrast, a mutant *C. trachomatis* enolase containing point substitutions at conserved residues at S44A lacked enzymatic activity (Figure 4B) (Brewer et al., 1998). These data demonstrate that *C. trachomatis* enolase is functional and verify that the activity that we measured was not due to contaminating *E. coli* enolase in our purified recombinant *C. trachomatis* enolase preparation.

**Figure 4.**
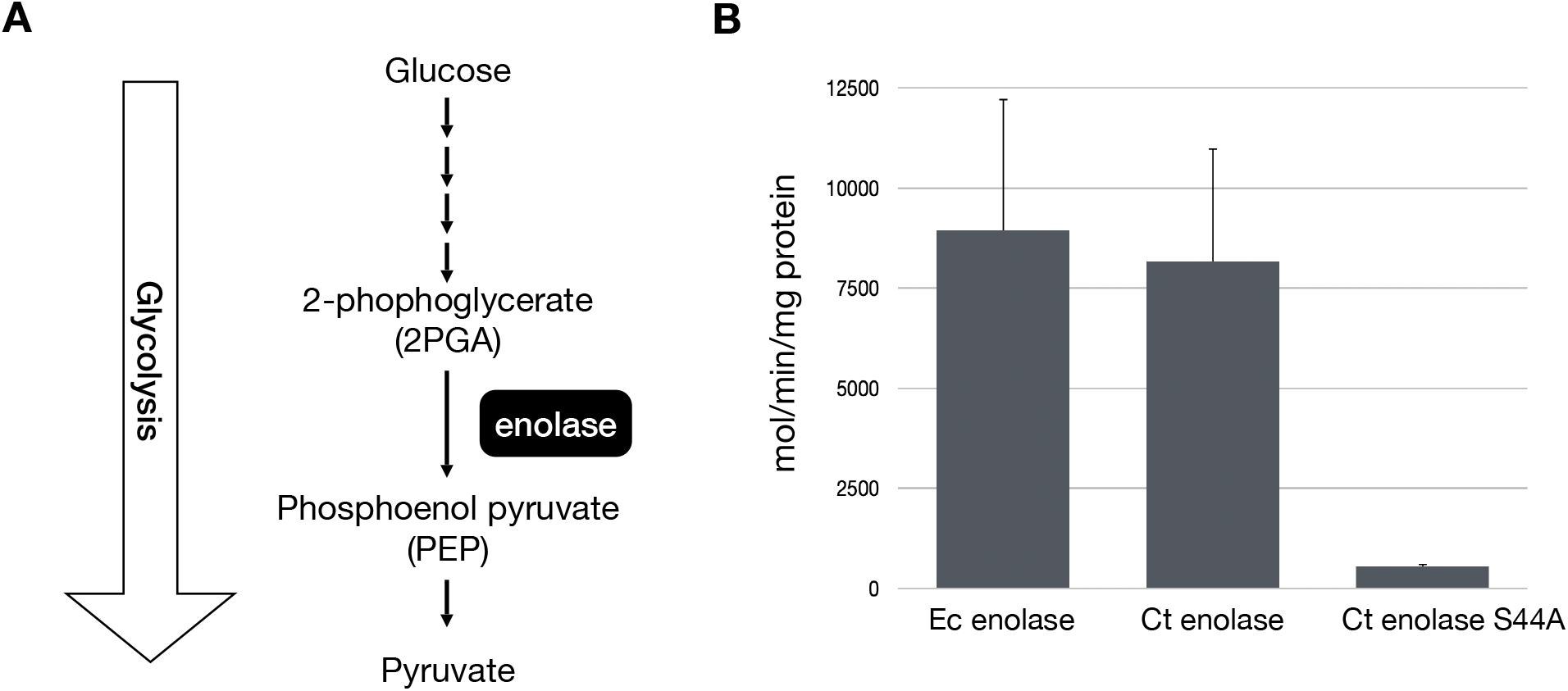
*C. trachomatis* enolase can convert 2PGA to PEP. A. Diagram of the glycolytic pathway and the enolase reaction step. B. In vitro enolase assay using purified recombinant *E. coli* enolase, *C. trachomatis* enolase, or the S44A mutant of *C. trachomatis* enolase. Enzymatic activity was calculated as the amount of PEP produced per minute per mg protein. Ec enolase is the *E. coli* recombinant protein Ct enolase and its mutant are *C. trachomatis* recombinant protein.

We next used our *in vitro* phosphatase assay to examine if enolase could be involved in regulating RsbU activity. In the presence of Mg^2+^ or high Mn^2+^, PEP inhibited the ability of RsbU to dephosphorylate RsbV1 and RsbV2 (Figure 5), while 2PGA did not. (Figure 5). We also tested TCA intermediates that have been shown to interact with the periplasmic domain of RsbU (Soules et al., 2020). 2-ketoglutarate and malic acid each had minimal effects on RsbU activity against RsbV1 and RsbV2, equivalent to the effect of the succinic acid negative control (Figure 6). Oxaloacetate (OAA) inhibited RsbU but only under specific conditions, i.e. against RsbV1 but not RsbV2, and only at low Mn^2+^ concentration, and not in the presence of high Mn^2+^ or Mg^2+^ concentration (Figure 6).

**Figure 5.**
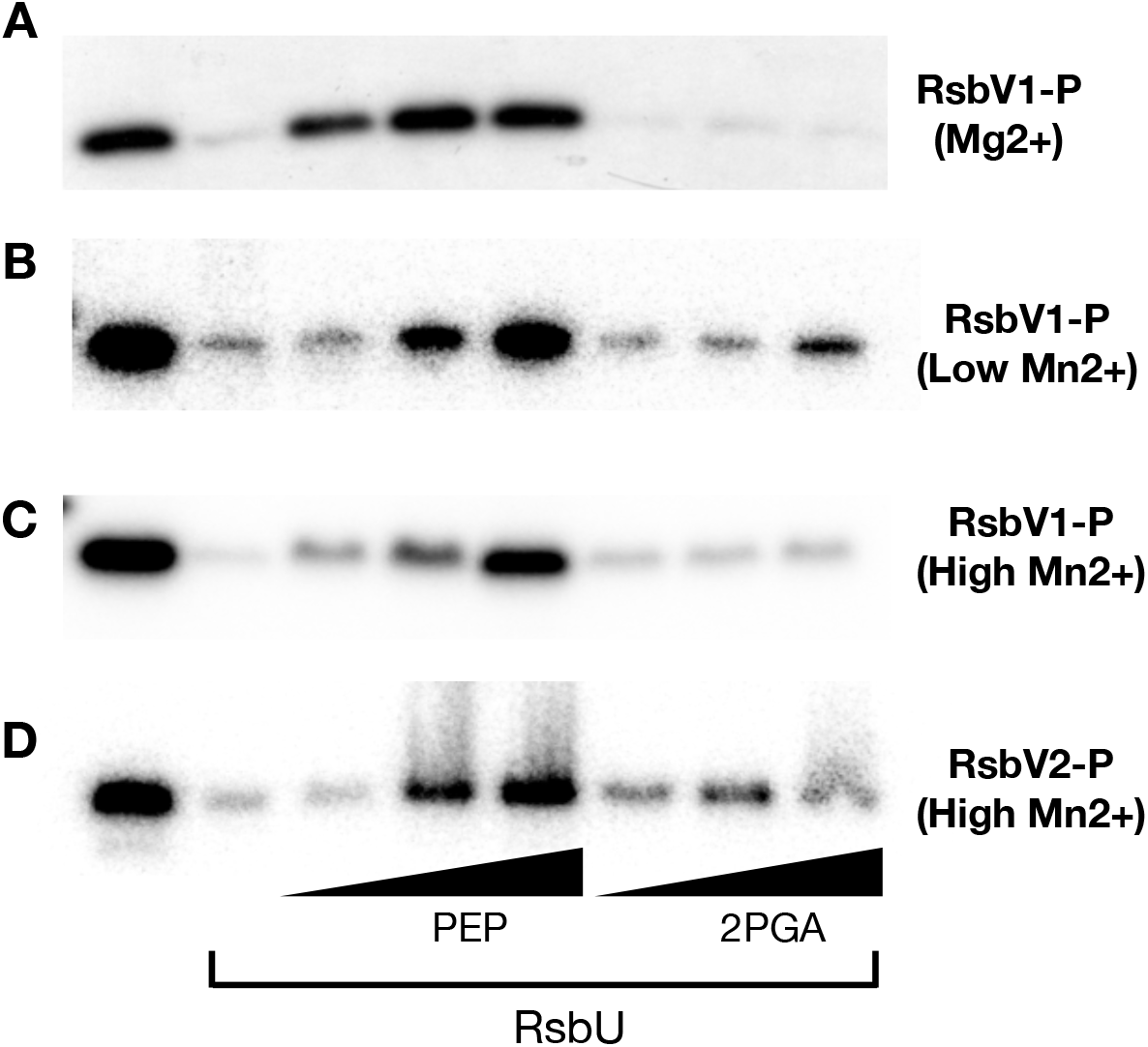
Inhibition of RsbU phosphatase activity by PEP. Autoradiographs showing *in vitro* phosphatase assay with purified recombinant RsbU (6 μM) and ^32^P-labelled RsbV1 and RsbV2 (5 μM) were performed with increasing concentrations (2.5, 5, and 10 mM) of PEP or 2PGA in the presence of (A) 10 mM MgCl_2_, (B) 0.1 mM MnCl_2_, (C) 10 mM MnCl_2_, or (D) 10 mM MnCl_2_ as the co-factor.

**Figure 6.**
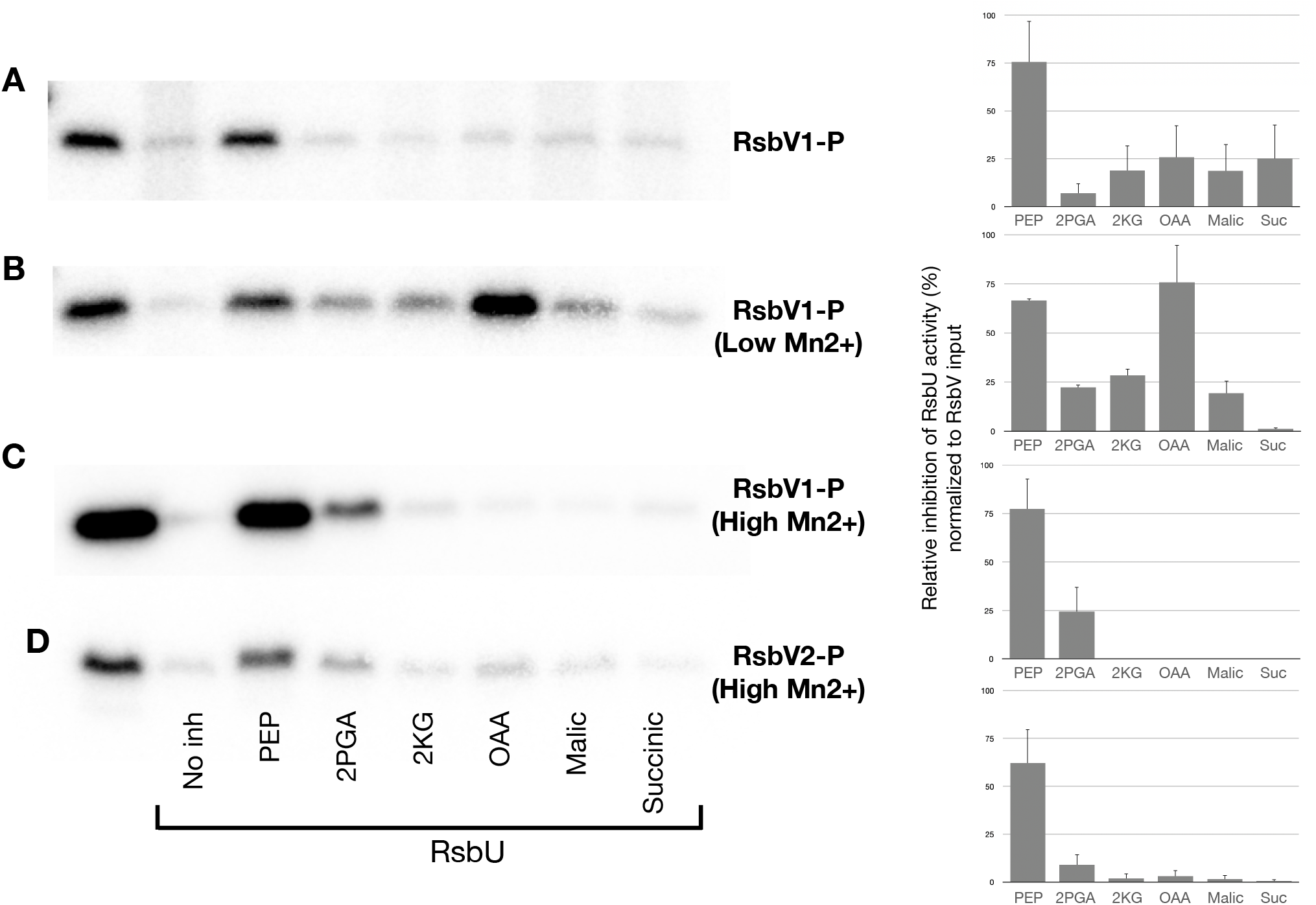
Effect of TCA intermediates on RsbU phosphatase activity. Autoradiographs showing *in vitro* phosphatase assay with purified recombinant RsbU (6 μM) and ^32^P-labelled RsbV1 and RsbV2 (5 μM) were performed with 10 mM of PEP, 2PGA, 2-ketoglutarate (2KG), oxaloacetate (OAA), malic acid, or succinic acid in the presence of (A) 10 mM MgCl_2_, (B) 0.1 mM MnCl_2_, (C) 10 mM MnCl_2_, and (D) 10 mM MnCl_2_ as the co-factor. Quantification of each autoradiograph shown to the right. Relative inhibition is expressed as the percent of inhibition, calculated as the amount of labelled RsbV divided by the RsbV input (these values were adjusted by subtracting the background of the no inhibitor control). Values represent the mean from at least three experiments with standard deviation indicated by the error bar.

## DISCUSSION

In this study, we characterized the phosphatase activity of *C. trachomatis* RsbU, including its co-factor requirement and putative substrates. We showed that manganese is a better co-factor than magnesium and that RsbU was able to dephosphorylate both RsbV1 and RsbV2. *C. trachomatis* RsbU has been previously shown to dephosphorylate RsbV1 (Thompson et al., 2015), but prior to this study it was not known if it could also dephosphorylate RsbV2. However, we found that RsbV2 was not as good a substrate as RsbV1, leaving open the question of whether RsbU or another enzyme is the cognate phosphatase for RsbV2 in *Chlamydia*.

The transcriptional organization of the *rsbU* gene in *C. trachomatis* provided a clue that enolase and RsbU might have a functional linkage in *Chlamydia*. The presence of *eno* and *rsbU* in the same operon was at first puzzling because there was no known relationship between glycolysis and the Rsb pathway of transcriptional regulation in other bacteria. The location of *eno* as the first gene in the *eno-rsbU* operon was particularly intriguing because the regulator of an operon is often its first gene, as is the case for the chlamydial transcription factors TrpR and HrcA (Akers and Tan, 2006; Hanson and Tan, 2015). This gene organization appears to be conserved, as *eno* is immediately upstream of *rsbU* in other *Chlamydia* spp., with the exception of *C. pneumoniae*, which has 5 genes between them.

Our data provide support for a novel functional connection between enolase and RsbU in *Chlamydia* through PEP, the product of the enolase reaction. We demonstrated that RsbU phosphatase activity is inhibited by PEP, but not by 2PGA, which is the substrate that is converted into PEP by enolase. These findings support a model in which enolase regulates RsbU by controlling the production of an inhibitor of this phosphatase.

Interestingly, RsbU may also be regulated by the TCA cycle. TCA intermediates, including 2-ketoglutarate, malic acid and oxaloacetate, have been reported to bind the C-terminal, periplasmic domain of RsbU (Soules et al., 2020). That study did not examine binding to the phosphatase domain. In our studies, 2-ketoglutarate and malic acid did not inhibit RsbU phosphatase activity, and oxaloacetate only inhibited RsbU activity under specific co-factor conditions and only against RsbV1 (Figure 6). It is possible that the periplasmic domain of RsbU is a regulatory domain and binding of TCA intermediates could inhibit RsbU phosphatase activity through allosteric mechanisms. We were unable to test this model because we could not purify full-length RsbU, and thus only performed experiments with the phosphatase domain of RsbU.

The ability of PEP to inhibit RsbU suggests several potential mechanisms for controlling the RsbW pathway and chlamydial gene expression. First, PEP production could be controlled by chlamydial glucose levels since PEP is a product of glucose metabolism through the glycolytic pathway. *Chlamydia*, as an intracellular bacterium, obtains glucose from the host cell in the form of glucose-6-phosphate as a carbon source (Schwoppe et al., 2002; Vender and Moulder, 1967; Weiss, 1965; Weiss and Wilson, 1969). Thus, enolase and RsbU could act as a sensor that activates transcription of Rsb-regulated genes when host glucose is limiting. Alternatively, PEP production could be controlled by enolase activity. In E. coli, enolase enzymatic activity is regulated by its phosphorylation state (Dannelly et al., 1989). Enolase may be developmentally regulated in Chlamydia because phosphorylated enolase has been detected in EBs but not in RBs in Chlamydia caviae (Fisher et al., 2015). Other potential regulators of enolase activity include inhibitors, such as fluoride, SF2312 phosphonate, and tropolone derivatives (Krucinska et al., 2019a; Krucinska et al., 2019b; Spring and Wold, 1971) or post-translational modification by lysine acetylation (Nakayasu et al., 2017).

We propose the following model for how enolase and RsbU could regulate gene expression in *Chlamydia* (Figure 7). When the host cell supplies sufficient glucose in the form of glucose-6-phosphate to RBs, the chlamydial glycolytic pathway produces 2PGA, which is converted into PEP by chlamydial enolase. PEP inhibits the phosphatase activity of RsbU, leaving its substrate RsbV in a phosphorylated form that is unable to bind RsbW. RsbW is thus free to bind its cognate sigma factor, inhibiting transcription by the form of RNA polymerase containing this sigma factor, be it sigma66 or sigma28. However, when host glucose is limiting, chlamydiae are unable to produce PEP, and RsbU becomes enzymatically active and dephosphorylates RsbV. Unphosphorylated RsbV binds RsbW, which frees up its cognate sigma factor so that it can direct the transcription of its target genes.This model provides a novel mechanism by which chlamydial gene expression could be regulated by nutrient availability.

**Figure 7.**
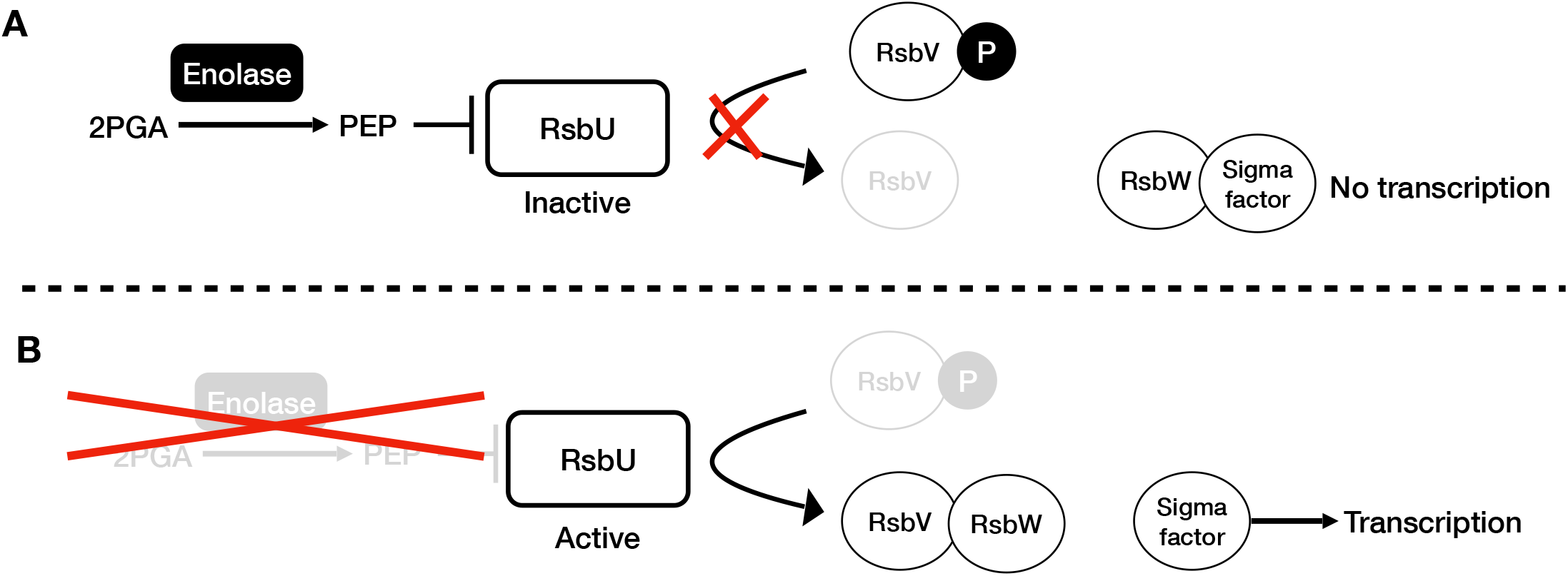
Model for regulation of RsbU phosphatase activity by enolase. A. PEP production by enolase inhibits transcription controlled by the Rsb pathway. PEP inhibits RsbU phosphatase activity, which results in accumulation of phosphorylated RsbV that is unable to bind RsbW. As a result.. RsbW binds to its cognate sigma factor and inhibit transcription of target genes. B. Active RsbU positively regulates transcription controlled by the Rsb pathway. In the absence of PEP, either because of decreased enolase activity or the lack of substrate (e.g. 2PGA or glucose) RsbU phosphatase is active and dephosphorylates RsbV. Unphosphorylated RsbV binds to RsbW, which frees up the cognate sigma factor to associate with other RNA polymerase subunits and transcribe target genes.

In summary, we have uncovered evidence of a functional connection between the glycolytic pathway and the Rsb pathway of gene regulation in *Chlamydia*. The linkage appears to be through enolase, which catalyzes the production of PEP, a novel inhibitor of chlamydial RsbU phosphatase activity. There has long been speculation about external stimuli that could regulate chlamydial gene expression and development, but upstream signals have not been identified (Chiarelli et al., 2020; Lee et al., 2018). Our findings provide a potential mechanism in which host glucose availability could be an upstream signal that regulates chlamydial gene expression.

## ACKNOWLEDGEMENTS

This work was supported by a grant from the NIH (AI044198).

---

| plasmid | Description |
| --- | --- |
| pMT1150 | *omcB* promoter |
| pMT1149 | *ompA* promoter |
| pMT1669 | *rsbU* promoter |
| pMT1781 | *E. coli* enolase in pRSET-C |
| pMT1673 | *C. trachomatis* enolase in pRSET-C |
| pMT1972 | *C. trachomatis* enolase containing a change at amino acid position 44 from a serine to an alanine in pRSET-C |
| pMT1135 | *C. trachomatis* RsbW in pRSET-C |
| pMT1136 | *C. trachomatis* RsbV2 in pRSET-C |
| pMT1754 | *C. trachomatis* RsbV1 in pRSET-C |
| pMT1676 | *C. trachomatis* RsbU containing amino acid 330-457 in pRSET-C |
| pRSET-C | Cloning plasmid (Invitrogen) |

